# *SCryPTA:* A web-based platform for analyzing Small-Angle Scattering curves of lyotropic liquid crystals

**DOI:** 10.1101/791848

**Authors:** Raphael Dias de Castro, Bruna Renata Casadei, Barbara Vasconcelos Santana, Mayra Lotierzo, Natália F. de Oliveira, Barbara Malheiros, Paolo Mariani, Renata C. K. Kaminski, Leandro R. S. Barbosa

## Abstract

Small angle X-ray scattering (SAXS) is a powerful technique for the characterization of systems with highly ordered structures, such as liquid crystals and self-assembly systems. In the field of nanotechnology and nanomedicine, SAXS can be used to characterize the crystallographic properties of the crystal phase of lyotropic systems and nanoparticles with internal crystal phase, such as cubosomes, hexosomes and multi-lamellar vesicles. In this work, we introduce a new web platform named *SCryPTA: Small Angle Scattering Crystallographic Peak Treatment and Analysis*, capable of reading SAXS data and providing a comprehensive visualization of the scattering curve along with the calculation of important physical parameters, such as the lattice parameter of the crystal structure, the lipidic bilayer width, among others. Cubic, hexagonal and multilamellar scattering data had their crystallographic structure characterized in SCryPTA. So far, four different cubic structures, (*Pn3m* (Q_224_), *Fd3m* (Q_227_), *Im3m* (Q_229_), *Ia3d* (Q_230_)), the hexagonal phase and also multi-lamellar vesicle systems are described in the platform. We believe that SCryPTA may help researchers from several fields, since it has a user-friendly interface. The platform is available at: www.if.usp.br/scrypta.

## INTRODUCTION

Amphiphilic molecules have a large spectra of applications, being used in basic sciences as well as in nanotechnology and nanomedicine^1,2^. Due to its molecular composition, they can self-assemble spontaneously in aqueous solution into several different lyotropic liquid crystalline (LLC) structures^3^. In particular, LLC structures can be mainly classified into different phases, as micellar, lamellar, cubic, and hexagonal according to their structures^4,5^. LLC are found in vivo^6,7^, they can be used in the crystallization of membrane proteins^8,9^, control of chemical reaction in food^10^ and as drug delivery systems^11–14^.

Small angle X-ray scattering (SAXS) is the most indicated technique to elucidate the polymorphism of LLC structure, since it can give information on the crystallographic structure of nanometric particles^15,16^. For the lamellar phase the water domains are localized between the bilayers, this is the typical structure of systems in the presence of a small amount of water^17^. The normal hexagonal phase (H_I_) can be understood as elongated (prolate) or infinite cylindrical micelles, being each one surrounded by six other micelles^4^. The inverse hexagonal phase (H_II_) consists of cylindrical inverted micelles of infinite length ordered into 2-D hexagonal arrangement^4^. Based on X-ray crystallographic studies, the cubic phase of lipids were described in the 60’s^18^. Later on, different cubic symmetries were elucidated, like the double-diamond like lattice (*Pn3m*), the body-centered lattice (*Im3m*), and the gyroid lattice cubic phase (*Ia3d*)^5,19^.

Despite the large amount of research dealing with LLC structures, it is not of our knowledge the existence of any web-based platform where the researcher can upload its scattering curve and get the main structural information of LLC-based systems. Thus, we report a web-based platform developed to study the structural features of LLC samples, analyzing the peaks in the SAXS curves. We show some examples of self-assembled structures that can be treated using SCryPTA.

SCryPTA can be used to describe cubic structures (*Pn3m, Ia3d, Im3m and Fd3m*), hexagonal systems and multilamellar vesicles. We provide a general overview of the platform, using some simple and complexes analysis, as follows.

## MATERIALS AND METHODS

### SAMPLES PREPARATION

The samples preparation and the experimental setup for SAXS measurements are described in the supplemental material.

#### SAXS analysis and computational tools

SCryPTA was developed using a Python-based interface, taking advantage of the features of the Jupyter platform. Thus, this web-based application uses well-known Python libraries, such as Numpy, Scipy, Matplotllib and Bokeh, and also a specific peak detection function also developed with Python.

The calculations performed to obtain the lattice parameter ***<u>a</u>*** and all the other derived parameters use a Gaussian fit procedure. Nevertheless, the user must know that this is an approach and it is quite often that diffraction peaks have no Gaussian profile. SCryPTA fits the data of the first diffraction peak, calculating structural parameter, depending on the symmetry.

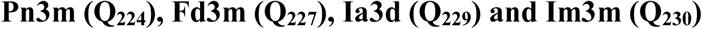

There are several studies in the literature describing cubic structures for lipids^5,9,20,21^. The structure of three of them (*Pn3m, Im3m, Ia3d*) can be described in terms of networks of cylinders, connected in different ways: in *Pn3m* the cylinders are tetrahedrally joined 4-by-4, similar to the diamond structure; for *Ia3d* the cylinders are coplanarly joined 3-by-3; and finally for *Im3m* the cylinders are cubically joined 6-by-6^5^. *Fd3m* structure, on the other hand, is composed by a series of micelles^5,22^.

SCryPTA can calculate structural parameters of cubic symmetries. For *Pn3m* and *Im3m*, SCryPTA can calculate the water channel (cylinder) radius (*r*_*w*_) and the length of the lipid molecule that forms the bilayer (*l*_*L*_), i.e, roughly half the length of the lipidic bilayer. In the case of the water channel radius, the following formulas reported in the literature were used^23,24^:

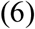

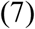

where *r*_*w*_ is the water channel radius, *a* is the lattice parameter of the respective cubic structure and *l*_*L*_ is the length of the lipid molecule in the bilayer, which in turn can be deduced from the lipid volume fraction *L* formula^4,25^:

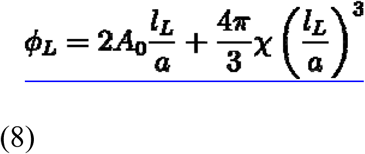

Equation 8 depends on two constant parameters related to the cubic phase: *A*_0_ is the ratio of the area of the minimal surface in the unit cell to the quantity ([*unit cell volume*]^2/3^) and is the Euler–Poincare characteristic number^24^, describing the topological space shape^4,25^. These values for the *Pn3m, Im3m* and *Ia3d* are presented in Table S1 in the supplemental material.

From Equation 8, *l*_*L*_ (Eq. 9) can be determined as^4^:

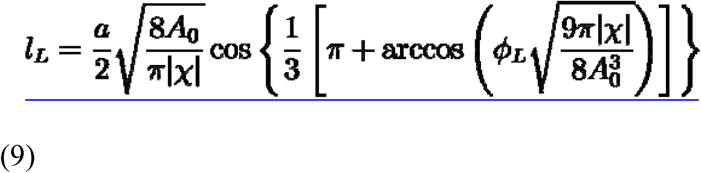

In addition to the cubic structures, SCryPTA can also calculate some important parameters for lamellar and hexagonal structures, such as the unit cell parameter.

#### Software Workflow

SCryPTA’s interface relies on the execution of several steps using the Jupyter notebook interface. Fig. 1A shows an overview of application’s workflow. Briefly, this tool can read and perform calculations from SAXS experimental data in which diffraction peaks are distinguishable from background scattering. The main function of this application is to identify the crystallographic symmetry present in the scattering pattern, and an optional peak detection function can be enabled to calculate some physical parameters, such as the lattice parameter.

**Figure 1.**
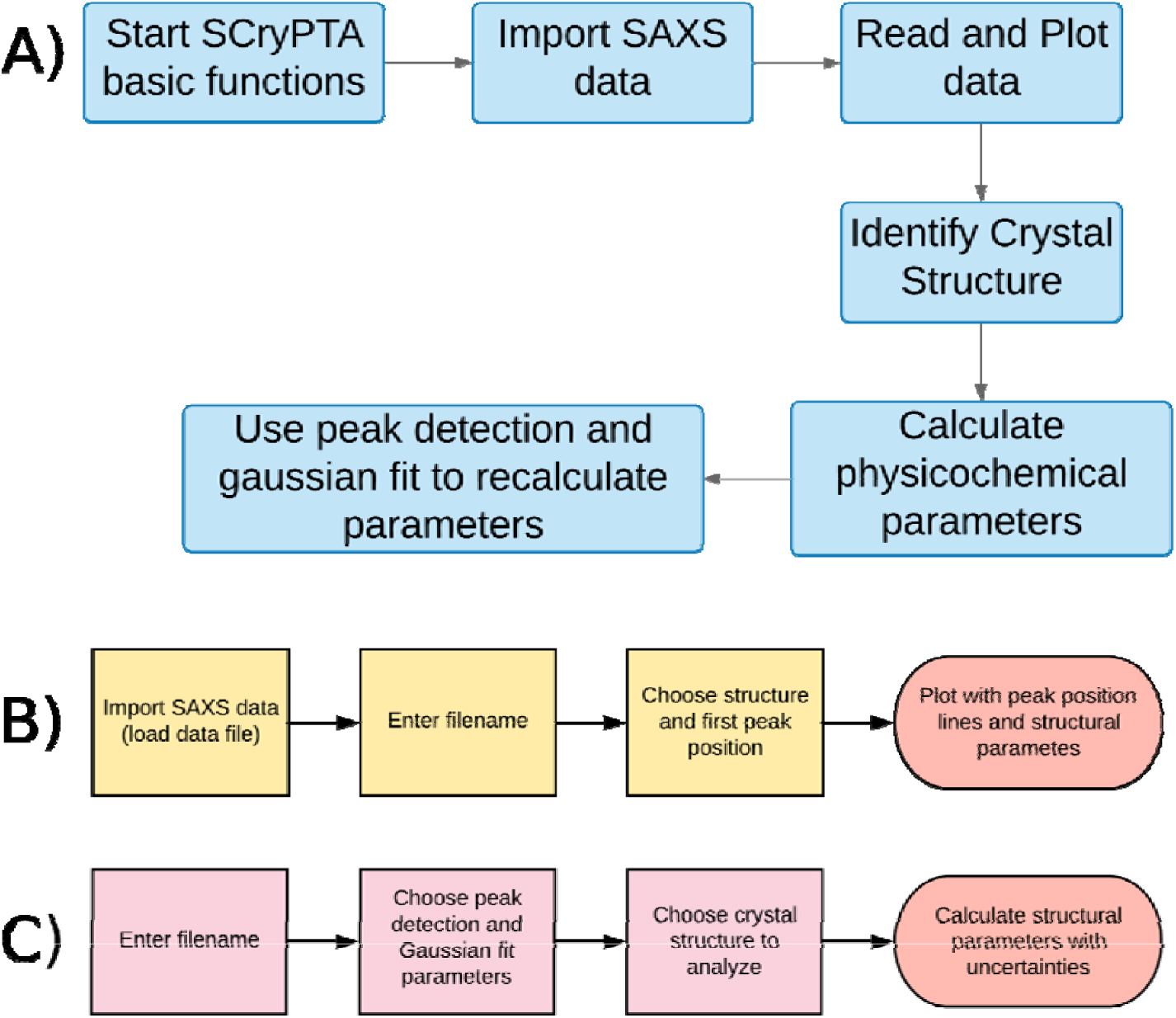
**(A)** SCryPTA flowchart with major processes in data analysis, composed of five parts: import SAXS data, read/plot data, identify crystal structure, use peak detection and gaussian fit to calculate parameters and plot data and show results. **(B)** Process for crystal lattice identification on SCryPTA has four steps: load data, enter filename with data to be analyzed, choose structure and first peak position, and finally generate plot. **(C)** An additional characterization process is provided in SCryPTA, where the user can calculate more accurate values for the structural parameters of the analyzed structure.

The most basic feature of SCryPTA is to provide easy and fast data visualization, which could be performed right after the data acquisition. Thus, using basic functions of Python’s Matplotlib and Bokeh libraries, SCryPTA produces an interactive plot that enables the researcher to explore the dataset.

In order to identify the crystal structure, SCryPTA provides an easy procedure (Fig. 1B). The user has the option to perform a visual “*initial guess*” for the position of the first peak and choose a specific symmetry available in the platform. SCryPTA will automatically plot lines indicating the expected position of the following 6 peaks, providing a visual fit of the expected diffraction peaks position. In addition to the identification of the crystal lattice, it also provides the calculations for the lattice parameter of the available structures, along with lipid length in the bilayer and water channel diameter for some cubic phases. These calculations are performed using the first peak position indicated by the user. This is a first approach to characterize the system related to the analyzed data (Fig. 1B).

SCryPTA can also provide a more accurate characterization using the implemented peak detection and fit functions (Fig. 1C). By automatically detecting the diffraction peaks present in the dataset, SCryPTA performs a gaussian fit to these peaks, generating the “fitted” position for each peak, with their respective uncertainties. These values are then used to calculate structural parameters for the selected structure.

The detailed use of SCryPTA can be visualized in the web-page manual, which is constantly updated as new versions of ScryPTA are released. (http://fig.if.usp.br/∼scrypta/Manual_SCryPTA.pdf). In the following section, we demonstrate how SCryPTA can be used to study LC systems, showing some examples.

## RESULTS AND DISCUSSION

In this section, some results obtained using SCryPTA are presented. Besides these examples, the reader can find more information on SCryPTA on the website (www.if.usp.br/scrypta). Initially, SCryPTA was implemented with four different cubic structures, besides the hexagonal and lamellar phases.

### 1. Cubic Structures

#### Monoolein

Monoolein (MO) is a nontoxic, biodegradable and biocompatible amphiphilic molecule used as emulsifying agent and food additive^26,27^. It exhibits a rich phase diagram with lamellar, cubic and hexagonal phases^5,25,28^. Figure 2A shows the SAXS data of MO in excess of water (25mg/ml) in the presence of Cytochrome-c (10mg/ml), along with two possible cubic structures: Im3m (red) and Pn3m (blue). As one can see, the Im3m symmetry is able to describe the peaks positions, whereas Pn3m is not, indicating that it is not appropriate to describe this system. SCryPTA also calculated the value of the lattice parameter 12.43±0.11 nm, which is in good agreement with the original results^5,12,25^. The lipidic bilayer length and the water channel diameter were also calculated by SCryPTA, being equal to 1.726±0.016 nm and 4.13±0.07 nm, respectively. Mariani et al.^5^ studied the structure of cubic phases of MO in the absence and presence of cytochrome-c, a water soluble protein found in the mitochondria^5^. Later on, Mazzoni et al.^25^ also studied the effect of cytochrome-c on MO phase diagram as a function of protein concentration, temperature and pressure^25^. The authors have shown that different concentrations of cytochrome-c in solution triggered a transition from *Pn3m* to *Im3m* (2 proteins per unit cell) cubic phases and from *Im3m* to *P4*_*3*_*32* cubic phase (10 to 20 proteins per unit cell)^25^. They also have shown that, at room temperature, the lattice parameter values were 10.60 nm for the *Pn3m* phase and 12.17 nm for the *Im3m* phase^25^, in good agreement with the values reported with SCryPTA.

**Figure 2:**
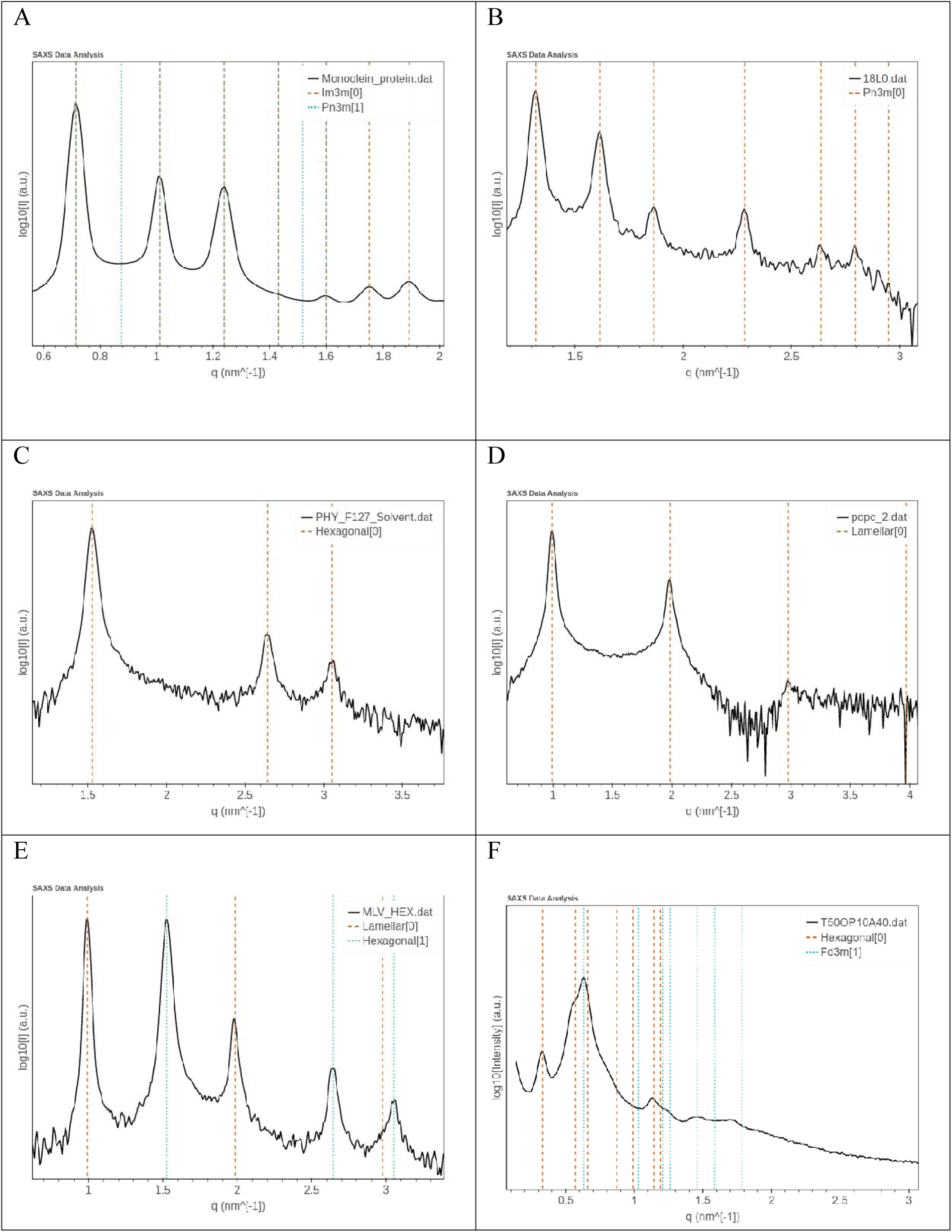
(**A**) Scattering pattern of the MO-water system in the presence of cytochrome-c, 10 mg/ml and MO at 25mg/ml^25^. SAXS curves of aqueous solution of cubosomes, composed of PHY and Pluronic F127 in the absence (B) and presence of 0.1 mM of chloroform (C). The red lines represent the theoretical peak positions of and hexagonal arrangement with inner lattice parameter of 4.755 ± 0.026 nm. (**D**) SAXS curve of MLVs composed by POPC at 10 mM in ultrapure milli-Q water. (**E**) Example of coexistence phase among hexagonal (green vertical lines) and MLVs (red vertical lines). (**F**) SAXS curve of the mixture: Tween80:Pracahy oil:water at 50:10:40 (%wt). Vertical lines indicate the presence of two structures: Hexagonal (red lines) and Fd3m (green lines).

#### Cubosomes (Pn3m)

Phytantriol (PHY) is a well-known active ingredient used in the cosmetic industry, such as hair and skin care products. PHY exhibits similar phase behavior to MO^29^ and has gained more recent interest in the biomedical field compared to monoglycerides due to several reasons, such as chemical stability due to the absence of the ester group and can be purchased commercially with high purity (∼95%)^20,29,30^.

Cubosomes are lipidic nanoparticles, with cubic inner structure^12,20,30^, they have shown interesting physicochemical properties and are being considered quite interesting as drug delivery systems^12,13,31^. They have higher hydrophilic surface area, being an interesting material for drug delivery systems^13,32^. The amphiphilic and non-ionic polymer Pluronic® F127 has a quite interesting behavior in the PHY physical properties. Under some specific conditions, stable cubosomes can be produced combining PHY and F127^30^.

Fig. 2B shows the SAXS curves of cubosomes composed of PHY and the amphiphilic polymer Pluronic® F127. As one can see, five-to-six peaks are present in the scattering pattern. Once the user defines the position of the first peak, SCryPTA can calculate the predicted positions of the next six peaks, according to the choose crystallographic symmetry. Fig. 2B shows the predicted peak positions for the *Pn3m* (vertical red lines) cubic phases. The first was defined at *q*_1_ = 1.302±0.008 nm^-1^. SCryPTA performed calculations of the structural parameters of this *Pn3m* structure. The lattice parameter was evaluated as 6.82±0.04 nm, in accordance with the results predicted in the literature ^49^. Also, two other important results were obtained: lipidic bilayer length at 1.412±0.009 nm and water channel diameter at 2.512±0.016 nm.

### 2. Hexagonal

Nonpolar organic solvents have been widely reported as a convenient method used to dissolve hydrophobic bioactive molecules^34–36^. Nevertheless, due to its hydrophobic nature, it may induce changes in nanoparticles depending on its concentration. Fig. 2C shows the SAXS diffraction pattern of PHY cubosomes in the presence of chloroform at 0.1 mM. A phase transition from cubic *Pn3m* to hexagonal (Fig. 2C) was verified using SCryPTA. The lattice parameter of the hexagonal phase was also provided by the application: 4.755±0.026 nm.

### 3. Multilamellar (MLVs)

Multilamellar vesicles (MLVs) are spherical concentric vesicles composed by several lamellar lipid bilayers. Fig. 2D shows the SAXS curve of POPC-MLVs. SCryPTA was able to identify the MLV phase in the sample, and also calculated the lattice parameter as 6.35±0.04 nm. Similar values are reported in the literature^37,38^. In this case, the lattice parameter is interpreted as the center-to-center average distance between two consecutive bilayers.

### 4. Phase Coexistence

SCryPTA was also developed to study phase coexistence. There are several cases where phase coexistence may occurs. Heftberger et al.^39^ studied the coexitence phase of Liquid ordered (L_o_) and Liquid disordered (L_d_) systems of lipids at increasing cholesterol concentrations. The authors found that cholesterol may reduce the difference between L_d_ and L_o_ thickness, decreasing the line tension and promoting the melting of L_o_ domain^39^. Figure 2E shows an example of lamellar (red lines) and hexagonal (blue lines) structures coexisting in solution. Scrypta is able to differentiate such structures, calculating the parameters of each one. Nevertheless, SCryPTA is not able to calculate the amount (percentage) of each structure in the sample.

Finally, SCryPTA can also be used to elucidate more complex structures. Pracachy oil is a not well studied cosmetic oil extracted from the seeds of *Pentaclethra macroloba* tree, which grows in Amazonian rainforest. It has interesting properties like antifungal, anti-inflammatory and anti-bacterial effects^40^. Besides that, it also presents capacity to heal scars fast^41^ and also has repellent effect against *Aedes aegypti*. TWEEN 80 is a well-known polysorbate frequently used in cosmetics and food. It has a well-studied structural behavior in aqueous solutions, forming micellar systems and gel-like LC structures, influenced by external factors such as pH, temperature, pressure and additives as oils and co-surfactants. Accordingly, all the scattering peaks can be represented as a sum of a cubic (*Fd3m*) with lattice parameter of 17.25 nm, and a hexagonal with unit cell of 21.99 nm (Fig. 2F). Surely, this is quite an unusual phase coexistence, and more measurements are being performed. Nevertheless, herein our goal is to show the capabilities of SCryPTA rather than the system itself.

## CONCLUSIONS

As we have shown along this manuscript, SCryPTA can successfully be used to describe the main structural features of LLC systems, with a user-friendly interface and calculating important structural features. So far, SCryPTA can be used to describe four different cubic structures, *Pn3m*(Q_224_), *Fd3m*(Q_227_), *Im3m*(Q_229_), *Ia3d*(Q_230_)) and both Hexagonal phases and Multilamellar stacking. In this version of SCryPTA it is also possible to elucidate a mixture of structures in the same sample given the structural parameters of each one (phase coexisting).. We believe that SCryPTA can be used by several different researchers, from nanostructured drug-delivery systems, including nanomedicine, engineering, food science, oil industry and technology. The platform is available at www.if.usp.br/scrypta.

## ACKNOWLEDGMENTS

Financial support for this research was provided by Fundação de Amparo à Pesquisa do Estado de São Paulo (FAPESP) (2015/15822-1, 2019/08832-1, 2018/04796-8), CAPES, CNPq (#155970/2018-6, 308692/2018-7, 420567/2016-0) and FAPITEC are also acknowledged. A special thanks to the National Laboratory of Synchrotron Light (LNLS), SAXS-1 beamline, Campinas/SP, for the usage of their facilities.

## Notes

http://fig.if.usp.br/~scrypta/

## REFERENCES

(1) Tadros, T. Self-Assembly of Surfactants. In Encyclopedia of Colloid and Interface Science; Tadros, T., Ed.; Springer Berlin Heidelberg: Berlin, Heidelberg, 2013, 1044–1044.

(2) Ranneh, A.-H.; Iwao, Y.; Noguchi, S.; Oka, T.; Itai, S. The Use of Surfactants to Enhance the Solubility and Stability of the Water-Insoluble Anticancer Drug SN38 into Liquid Crystalline Phase Nanoparticles. Int. J. Pharm. 2016, 515, 501–505.

(3) Israelachvili, J. N. Intermolecular and Surface Forces: Revised Third Edition. Acad. Press. 2011.

(4) Marsh, D. Handbook of Lipid Bilayers, 2nd ed.; CRC Press: Boca Raton, FL, USA, 2013.

(5) Mariani, P.; Luzzati, V.; Delacroix, H. Cubic Phases of Lipid-Containing Systems. Structure Analysis and Biological Implications. J. Mol. Biol. 1988, 204, 165–189.

(6) Wilts, B. D.; Apeleo Zubiri, B.; Klatt, M. A.; Butz, B.; Fischer, M. G.; Kelly, S. T.; Spiecker, E.; Steiner, U.; Schröder-Turk, G. E. Butterfly Gyroid Nanostructures as a Time-Frozen Glimpse of Intracellular Membrane Development. Sci. Adv. 2017, 3, e1603119.

(7) Almsherqi, Z. A.; Landh, T.; Kohlwein, S. D.; Deng, Y. Chapter 6: Cubic Membranes the Missing Dimension of Cell Membrane Organization. Int. Rev. Cell Mol. Biol. 2009, 274, 275–342.

(8) Zabara, A.; Chong, J. T. Y.; Martiel, I.; Stark, L.; Cromer, B. A.; Speziale, C.; Drummond, C. J.; Mezzenga, R. Design of Ultra-Swollen Lipidic Mesophases for the Crystallization of Membrane Proteins with Large Extracellular Domains. Nat. Commun. 2018, 9, 1–9.

(9) Landau, E. M.; Rosenbusch, J. P. Lipidic Cubic Phases: A Novel Concept for the Crystallization of Membrane Proteins. Proc. Natl. Acad. Sci. U. S. A. 1996, 93, 14532–14535.

(10) Mezzenga, R.; Schurtenberger, P.; Burbidge, A.; Michel, M. Understanding Foods as Soft Materials. Nat. Mater. 2005, 4, 729–740.

(11) Radaic, A.; Pugliese, G. O.; Campese, G. C.; Pessine, F. B. T.; Jesus, M. B. de; Radaic, A.; Pugliese, G. O.; Campese, G. C.; Pessine, F. B. T.; Jesus, M. B. de. Como estudar interações entre nanopartículas e sistemas biológicos. Quím. Nova 2016, 39, 1236–1244.

(12) Carducci, F.; Casadei, B. R.; Mariani, P.; Barbosa, L. R. S. X-Ray Characterization of Pharmaceutical and Cosmetic Lipidic Nanoparticles for Cutaneous Application. Curr. Pharm. Des. 2019, 25, 2364–2374.

(13) Esposito, E.; Carducci, F.; Mariani, P.; Huang, N.; Simelière, F.; Cortesi, R.; Romeo, G.; Puglia, C. Monoolein Liquid Crystalline Phases for Topical Delivery of Crocetin. Colloids Surf. B Biointerfaces 2018, 171, 67–74.

(14) Esposito, E.; Drechsler, M.; Puglia, C.; Cortesi, R. New Strategies for the Delivery of Some Natural Anti-Oxidants with Therapeutic Properties. Mini Rev. Med. Chem. 2019, 19, 1030-1039.

(15) Wang, H.; Zetterlund, P. B.; Boyer, C.; Spicer, P. T. Polymerization of Cubosome and Hexosome Templates to Produce Complex Microparticle Shapes. J. Colloid Interface Sci. 2019, 546, 240–250.

(16) Lv, F.; An, Z.; Wu, P. Scalable Preparation of Alternating Block Copolymer Particles with Inverse Bicontinuous Mesophases. Nat. Commun. 2019, 10:1397.

(17) Zhai, J.; Fong, C.; Tran, N.; Drummond, C. J. Non-Lamellar Lyotropic Liquid Crystalline Lipid Nanoparticles for the Next Generation of Nanomedicine. ACS Nano 2019, 13, 6178–6206.

(18) Luzzati, V.; Spegt, P. A. Polymorphism of Lipids. Nature 1967, 215, 701–704.

(19) Wang, X.; Zhang, Y.; Gui, S.; Huang, J.; Cao, J.; Li, Z.; Li, Q.; Chu, X. Characterization of Lipid-Based Lyotropic Liquid Crystal and Effects of Guest Molecules on Its Microstructure: A Systematic Review. AAPS PharmSciTech 2018, 19, 2023–2040.

(20) Rizwan, S. B.; Dong, Y.-D.; Boyd, B. J.; Rades, T.; Hook, S. Characterisation of Bicontinuous Cubic Liquid Crystalline Systems of Phytantriol and Water Using Cryo Field Emission Scanning Electron Microscopy (Cryo FESEM). Micron Oxf. Engl. 1993 2007, 38, 478–485.

(21) Chountoulesi, M.; Pippa, N.; Pispas, S.; Chrysina, E. D.; Forys, A.; Trzebicka, B.; Demetzos, C. Cubic Lyotropic Liquid Crystals as Drug Delivery Carriers: Physicochemical and Morphological Studies. Int. J. Pharm. 2018, 550, 57–70.

(22) Rappolt, M.; Cacho-Nerin, F.; Morello, C.; Yaghmur, A. How the Chain Configuration Governs the Packing of Inverted Micelles in the Cubic Fd 3 M-Phase. Soft Matter 2013, 9, 6291–6300.

(23) Briggs, J.; Caffrey, M. The Temperature-Composition Phase Diagram and Mesophase Structure Characterization of Monopentadecenoin in Water. Biophys. J. 1994, 67, 1594–1602.

(24) Tyler, A. I. I.; Barriga, H. M. G.; Parsons, E. S.; McCarthy, N. L. C.; Ces, O.; Law, R. V.; Seddon, J. M.; Brooks, N. J. Electrostatic Swelling of Bicontinuous Cubic Lipid Phases. Soft Matter 2015, 11, 3279–3286.

(25) Mazzoni, S.; Barbosa, L. R. S.; Funari, S. S.; Itri, R.; Mariani, P. Cytochrome-c Affects the Monoolein Polymorphism: Consequences for Stability and Loading Efficiency of Drug Delivery Systems. Langmuir 2016, 32, 873–881.

(26) Bozzola, R. Monoolein and emulsions for internal use. Boll Chim Farm 1954, 93, 415–417.

(27) Strauss, E. W. Electron Microscopic Study of Intestinal Fat Absorption in Vitro from Mixed Micelles Containing Linolenic Acid, Monoolein, and Bile Salt. J. Lipid Res. 1966, 7, 307–323.

(28) Qiu, H.; Caffrey, M. The Phase Diagram of the Monoolein/Water System: Metastability and Equilibrium Aspects. Biomaterials 2000, 21, 223–234.

(29) Barauskas, J.; Landh, T. Phase Behavior of the Phytantriol/Water System. Langmuir 2003, 19, 9562–9565.

(30) Akbar, S.; Anwar, A.; Ayish, A.; Elliott, J. M.; Squires, A. M. Phytantriol Based Smart Nano-Carriers for Drug Delivery Applications. Eur. J. Pharm. Sci. 2017, 101, 31–42.

(31) Barriga, H. M. G.; Holme, M. N.; Stevens, M. M. Cubosomes; the next Generation of Smart Lipid Nanoparticles? Angew. Chem. Int. Ed Engl. 2019, 10, 2958–2978.

(32) Duttagupta, A. S.; Chaudhary, H. M.; Jadhav, K. R.; Kadam, V. J. Cubosomes: Innovative Nanostructures for Drug Delivery. Curr. Drug Deliv. 2016, 13, 482–493.

(33) Akhlaghi, S. P.; Ribeiro, I. R.; Boyd, B. J.; Loh, W. Impact of Preparation Method and Variables on the Internal Structure, Morphology, and Presence of Liposomes in Phytantriol-Pluronic(®) F127 Cubosomes. Colloids Surf. B Biointerfaces 2016, 145, 845–853.

(34) Chen, J.-Y.; Su, C.-Y.; Hsu, C.-H.; Zhang, Y.-H.; Zhang, Q.-C.; Chang, C.-L.; Hua, C.-C.; Chen, W.-C. Solvent Effects on Morphology and Electrical Properties of Poly(3-Hexylthiophene) Electrospun Nanofibers. Polymers 2019, 11, 1501.

(35) Magiera, A.; Marchelak, A.; Michel, P.; Owczarek, A.; Olszewska, M. A. Lipophilic Extracts from Leaves, Inflorescences and Fruits of Prunus Padus L. as Potential Sources of Corosolic, Ursolic and Oleanolic Acids with Anti-Inflammatory Activity. Nat. Prod. Res. 2019, 1–6.

(36) Meli, V.; Caltagirone, C.; Falchi, A. M.; Hyde, S. T.; Lippolis, V.; Monduzzi, M.; Obiols-Rabasa, M.; Rosa, A.; Schmidt, J.; Talmon, Y.; et al. Docetaxel-Loaded Fluorescent Liquid-Crystalline Nanoparticles for Cancer Theranostics. Langmuir ACS J. Surf. Colloids 2015, 31, 9566–9575.

(37) Boulgaropoulos, B.; Rappolt, M.; Sartori, B.; Amenitsch, H.; Pabst, G. Lipid Sorting by Ceramide and the Consequences for Membrane Proteins. Biophys. J. 2012, 102, 2031–2038.

(38) Heftberger, P.; Kollmitzer, B.; Heberle, F. A.; Pan, J.; Rappolt, M.; Amenitsch, H.; Kucerka, N.; Katsaras, J.; Pabst, G. Global Small-Angle X-Ray Scattering Data Analysis for Multilamellar Vesicles: The Evolution of the Scattering Density Profile Model. J. Appl. Crystallogr. 2014, 47 (1), 173–180.

(39) Heftberger, P.; Kollmitzer, B.; Rieder, A. A.; Amenitsch, H.; Pabst, G. In Situ Determination of Structure and Fluctuations of Coexisting Fluid Membrane Domains. Biophys. J. 2015, 108, 854–862.

(40) Santiago, G. M. P.; Viana, F. A.; Pessoa, O. D. L.; Santos, R. P.; Pouliquen, Y. B. M.; Arriaga, A. M. C.; Andrade-Neto, M.; Braz-Filho, R. Avaliação Da Atividade Larvicida de Saponinas Triterpênicas Isoladas de Pentaclethra Macroloba (Willd.) Kuntze (Fabaceae) e Cordia Piauhiensis Fresen (Boraginaceae) Sobre Aedes Aegypti. Rev. Bras. Farmacogn. 2005, 15, 187–190.

(41) Banov, D.; Banov, F.; Bassani, A. S. Case Series: The Effectiveness of Fatty Acids from Pracaxi Oil in a Topical Silicone Base for Scar and Wound Therapy. Dermatol. Ther. 2014, 4, 259–269.

