## Supplemental material for "*SCryPTA:* A web-based platform for analyzing Small-Angle Scattering curves of lyotropic liquid crystals"

**MATERIALS AND METHODS**

**SAMPLES PREPARATION**

*Fully Hydrated Monoolein:* fully hydrated lipid samples were prepared by soaking monoolein (MO, glyceryl monooleate or 2,3-dihydroxypropyl oleate, with >99%) into the appropriate amount of ultrapure water in the absence and presence of cytochrome-c, a water-soluble protein present in the mitochondria^1,2^

. MO final concentration was set to c_MO_=25 mg/mL. After preparation, MO samples were vortexed and left for equilibration for 4 hours before the measurements.

*Cubosomes* and *Hexosomes:* samples prepared with phytantriol (PHY, 3,7,11,15- tetramethyl-1,2,3-hexadecanetriol) and Pluronic F-127 were based on bottom-up methodology, studied by Akhlaghi et al.^3^. Briefly, two solutions were produced, the first one 100 mg of PHY was solubilized into 10 mL of ethanol. Then, 25 mg of F127 was solubilized in 22.5 mL of ultrapure water. Subsequently, lipid solution was dropwise into the F127, at 45 ºC (above fusion point). The final solution was brought to a rotary evaporator until the final volume was 5 mL. In order to produce the hexosomes, 0.1 mM of chloroform was added in the regular cubosome sample.

*Liposomes:* samples were prepared by solubilizing the 10 mM of POPC (1-palmitoyl-2-oleoyl-*sn*-glycero-3-phosphocholine) in chloroform. The solvent was evaporated with a continuous flow of ultra-pure N_2_ forming a lipid film, and then, stored at low pressure (inside desiccator, using a vacuum pump) for 2 hours. Ultrapure water was added to the test tube, and multilamellar vesicles (MLVs) were formed by mechanical agitation of the tube.

*Amazonian rainforest oils:* liquid-crystalline formulations based in Amazonian rainforest oil were prepared as: Polysorbate 80 (Tween 80), *Pentaclethra macroloba* oil (Pracachy oil, PO) and ultrapure water at 50:10:40 (%wt), respectively. All components were mixed manually. Formulations were stabilized at room temperature by 18 hours and stored at 15°C.

**Small Angle X-ray Scattering**

SAXS experiments were performed at beamline D01B, SAXS1 at the Brazilian Synchrotron Light Laboratory (LNLS), Campinas, Brazil. Beam wavelength was 1.488 Å and the sample-detector distance ~1000 mm, the time acquisition was set to 100 s and all measurements were performed at room temperature (22.0±0.5 ^o^C). SAXS curves were obtained by radially average the 2D images of the detector (300K Pilatus detector). The contribution of the buffer solution was subtracted considering the sample's attenuation.

Table S1 describes the values of the parameter used in Eq. 8 and Eq. 9.

Table S1 – Values for the cubic minimal surface parameters^2^.

| ***Cubic phase*** | ***A_0_*** | ***𝜒*** |
| --- | --- | --- |
| *Pn3m* | 1.919 | -2 |
| *Im3m* | 2.345 | -4 |
| *Ia3d* | 3.091 | -8 |

**REFERENCES**

(1)	Mariani, P.; Luzzati, V.; Delacroix, H. Cubic Phases of Lipid-Containing Systems. Structure Analysis and Biological Implications. *J. Mol. Biol.* **1988**, *204*, 165–189.

(2)	Mazzoni, S.; Barbosa, L. R. S.; Funari, S. S.; Itri, R.; Mariani, P. Cytochrome-c Affects the Monoolein Polymorphism: Consequences for Stability and Loading Efficiency of Drug Delivery Systems. *Langmuir* **2016**, *32*, 873–881.

(3)	Akhlaghi, S. P.; Ribeiro, I. R.; Boyd, B. J.; Loh, W. Impact of Preparation Method and Variables on the Internal Structure, Morphology, and Presence of Liposomes in Phytantriol-Pluronic(®) F127 Cubosomes. *Colloids Surf. B Biointerfaces* **2016**, *145*, 845–853.
